# Genome annotation of Poly(lactic acid) degrading *Pseudomonas aeruginosa* and *Sphingobacterium sp.*

**DOI:** 10.1101/609883

**Authors:** Sadia Mehmood Satti, Aamer Ali Shah, Rafael Auras, Terence L. Marsh

## Abstract

*Pseudomonas aeruginosa* and *Sphinogobacterium sp*. are well known for their ability to decontaminate many environmental pollutants like PAHs, dyes, pesticides and plastics. The present study reports the annotation of genomes from *P. aeruginosa* and *Sphinogobacterium sp*. that were isolated from compost, based on their ability to degrade poly(lactic acid), PLA, at mesophillic temperatures (~30°C). Draft genomes of both the strains were assembled from Illumina reads, annotated and viewed with an aim of gaining insight into the genetic elements involved in degradation of PLA. The draft-assembled genome of strain *Sphinogobacterium* strain S2 was 5,604,691 bp in length with 435 contigs (maximum length of 434,971 bp) and an average G+C content of 43.5%. The assembled genome of *P. aeruginosa* strain S3 was 6,631,638 bp long with 303 contigs (maximum contig length of 659,181 bp) and an average G+C content 66.17 %. A total of 5,385 (60% with annotation) and 6,437 (80% with annotation) protein-coding genes were predicted for strains S2 and S3 respectively. Catabolic genes for biodegradation of xenobiotic and aromatic compounds were identified on both draft genomes. Both strains were found to have the genes attributable to the establishment and regulation of biofilm, with more extensive annotation for this in S3. The genome of *P. aeruginosa* S3 had the complete cascade of genes involved in the transport and utilization of lactate while *Sphinogobacterium strain* S2 lacked lactate permease, consistent with its inability to grow on lactate. As a whole, our results reveal and predict the genetic elements providing both strains with the ability to degrade PLA at mesophilic temperature.

## 1. Introduction

Poly(lactic acid) (PLA), is a bio-based aliphatic polyester polymer, obtained from sources such as corn sugar, cassava, wheat, rice, potato, and sugar cane, considered renewable [1, 2]. PLA is completely biodegradable under industrial composting conditions [3] as well as under unsupervised environmental conditions where its biodegradation is considered safe [4]. In the last two decades biodegradation of PLA has been extensively studied and many microbial species (actinomycete, bacteria, fungus) have been identified with the ability to degrade PLA [4]. Most of the reported bacterial species are from the family *Pseudonocardiaceae, Thermomonosporaceae*, *Micromonosporaceae*, *Streptosporangiaceae*, *Bacillaceae* and *Thermoactinomycetaceae* while the fungal species are mainly from the phylum *Basidiomycota* (*Tremellaceae*) and *Ascomycota* (*Trichocomaceae*, *Hypocreaceae*) [5–11].

In our previous study we also described four bacterial strains designated as S1, S2, S3 and S4, able to degrade PLA at ambient temperature [12]. Two of the isolated strains, *Sphingobacterium sp*. (S2) and *P. aeruginosa (S3)*, were also evaluated for their PLA degradation in soil microcosms [13]. The genus *Sphingobacterium* is from Phylum Bacteriodetes, Family *Sphingobacteriace*, named with reference to the sphingolipids in their cell wall [14, 15]. They are gram-negative rods and the GC content of their DNA is usually ranging from 35 to 44 mol% [16, 17]. *Sphingobacterium sp*. are found in a range of habitats like soil, forest, compost, activated sludge, rhizosphere, faeces, lakes and various food sources [18]. *Sphingobacterium* had also been reported to have their potential role in biodegradation of different pollutants including mixed plastic waste, PAHs, biodegradation of oil and pesticides [19–21]. *Pseudomonas aeruginosa* are gram-negative bacteria from γ-subdivision of proteobacteria. They are ubiquitously distributed in soil and aquatic habitats and are well-known opportunistic pathogens [22, 23]. It has the ability to thrive in highly diverse and unusual ecological niches with scarce available nutrients. Its metabolic versatility allows it to survive on a variety of diverse carbon sources for its survival, even in some disinfectants. and can metabolize many antibiotics [24, 25]. In previous reports role of *Pseudomonas aeruginosa* in degradation of different polymers including PAHs, biodegradation of xenobiotic compounds, degradation of oil, dyes and plastics is well documented [26–30],

PLA degrading bacteria reported in our previous study [12] were isolated from compost and had the ability to degrade PLA at ambient temperature. Interestingly, our *P. aeruginosa* strains were lactate utilizing while the *Sphingobacterium sp*. and *Chryseobacterium sp*. strains were unable to utilize lactate when provided as the sole carbon source in minimal media. We also observed that all four isolates could form biofilm on PLA. The purpose of this study was to analyze the genomes of two of our isolates and explore the genetic determinants responsible for conferring the particular characteristics to the strains promoting their degradation ability. The genes controlling lactate utilization and biofilm formation and regulation were identified. Whole genome sequence analysis for *P. aeruginosa* is extensive but such data for *Sphingobacterium sp*. is quite limited. To our knowledge this is first report that gives such genetic depth to PLA degrading bacterial strains and will provide the basis for further analysis.

## 2. Material and methods

### 2.1 DNA extraction

Two of our previously isolated, PLA degrading bacterial strains, *Sphingobacterium* sp. strain S2 and *P. aeruginosa* strain S3 (GenBank accession numbers *KY432687* and *KY432688*, respectively) were selected for genome sequencing [12]. Both of these strains were grown separately in 100 mL of LB in a 250 mL Erlenmeyer flask for 16 hours in a shaking incubator at 30°C and 70 rpm. Genomic DNA was subsequently isolated by using MO BIO PowerSoil^®^ DNA isolation kit (MO BIO laboratories, Inc. Loker Ave west, Carlsbad, CA). NanoDrop^®^ ND-1000 spectrophotometer and ND-1000 V3.1.8 software (Wilmington, DE, USA) was used to determine DNA concentrations of purified samples and sent for whole genome sequencing at Michigan State University Genomics Facility (MSU-RTSF).

### 2.2 Genome sequencing

Libraries for sequencing were prepared using the Illumina TruSeq Nano DNA Library Preparation Kit on a Perkin Elmer Sciclone NGS robot. Before sequencing, the qualities of the libraries were tested and quantification was performed using a combination of Qubit dsDNA HS, Caliper LabChip GX HS DNA and Kapa Illumina Library Quantification qPCR assays. Libraries were pooled in equimolar quantities and loaded on an Illumina MiSeq standard v2 flow cell with a 2×250bp paired end format and using a v2 500 cycle reagent cartridge. Illumina Real Time Analysis (v1.18.64) was used for base calling and the output was converted to FastQ format with Illumina Bc12fastq (v1.8.4) after demultiplexing. A total of 6,304,420 reads (~3.15 GB) were obtained for strain S2 and 5,800,229 reads (~2.9GB) were obtained for strain S3.

### 2.3 Sequence assembly, annotation and analysis

Assembly of the whole genome was performed using the full Spades assembly function within PATRIC (Pathosystems Resource Integration Center) (PATRIC 3.4.9) as implemented in the *miseq* assembly option. This assembly option incorporates BayesHammer algorithms followed by Spades, (Spades version 3.8.). Rast tool kit as implemented in PATRIC (PATRIC 3.4.9) was used for the annotation of contigs. The assembled contig file generated from this assembly was used as seed for the Comprehensive Genome Analysis function in PATRIC. The genomes were interrogated for the distribution of specific protein families (PGFams) using the protein family sorter tool on PATRIC. The genomes were compared to their closest reference genomes available on PATRIC to examine the strain-specific unique proteins as well as proteins common to the closest relative using the filter option in protein family sorter tool on PATRIC.

### 2.4 Average nucleotide identity (ANI) for species delineation

Isolates were further analyzed using a whole genome based Average nucleotide identity (ANI) method to delineate the genomes to their correctly. ANI values were calculated using MiSI (microbial species identifier) tool that is publicly available at Integrated Microbial Genomes (IMG) database [31]. The algorithm used in the original method proposed by Konstantinidis and Tiedje was modified and used to determine ANI between two genomes [32]. The average of the nucleotide identity of the orthologous genes of the pair of genomes was calculated and identified as bidirectional best hits (BBHs) using a similarity search tool, NSimScan (http://www.scidm.org/). The ANI of one genome to other genome is defined as the sum of the %-identity times the alignment length for all best bidirectional hits, divided by the sum of the lengths of the BBH genes. This pairwise calculation is performed in both directions. The strains used for comparison were complete genomes obtained from NCBI and are as follows. For the *Sphingobacterium* comparisons; *S. thalpophilum* DSM 11723 (Draft genome of 32 contigs), *S. sp*. G1-14, S. sp. B29, *S. multivorum* DSM 11691, *S. lactis* DSM 22361, *S. wenxiniae* DSM 22789, *S. mizutaii* DSM 11724 and *S. sp*. 21. For the *Pseudomonas aeruginosa* comparisons; *P. aeruginosa* PSE302, *P. aeruginosa* PA96, *P. aeruginosa* PA01H20, *P. aeruginosa* DSM 50071 *P. aeruginosa* PAO1, *P. aeruginosa* PAK, *P. aeruginosa* O12 PA7, *P. aeruginosa* PA_D25, *P. aeruginosa* PA_D1, *P. aeruginosa* KU and *P. aeruginosa* T52373.

### 2.5 Comparative alignments using MeDuSa and Mauve

For the comparative alignment of the genomes with their reference genomes and their visualization, MeDuSa [33] was used to reduce the number of contigs through comparison with the gene order of the closest strain. This was followed by alignment with MAUVE [34] to the same reference strain to provide an estimate of alignment similarity. *P. aeruginosa* PSE305 was used as a reference for our *P. aeruginosa* S3 while Sphingobacterium thalpophilum DSM 11723 was selected as a reference for *Sphingobacterium sp*. S2. References were selected based on closest ANI score.

## 3. Results and discussion

### 3.1. General Genome features of *Sphingobacterium* sp. (S2) and *P. aeruginosa* (S3)

The assembly of the draft genome for strain S2 yielded 87 contigs (434971 maximum length) and a total of 5,445,390 assembled base pairs with 43.66% G+C (Table 1). The largest contig was 434,944 bp. There were an estimated 4,951 CDS regions, 2,864 proteins with functional assignments and several antibiotic resistance genes. The assembled draft genome of S3 had 63 contigs (658,980 maximum length) with a total of 6,509,961 assembled base pairs and was 66.26% G+C. There were an estimated 6,239 CDS regions and 4,932 proteins with functional assignments and a substantial number of putative antibiotic resistant and virulence determinants.

**Table 1.** General genomic features of *Sphingobacterium* sp. S2 and *P. aeruginosa* S3

| Features | *Sphingobacterium* S2 | *Pseudomonas* S3 | Database<sup>1</sup> |
| --- | --- | --- | --- |
| Contigs | 87 | 63 | PATRIC |
| GC Content | 43.66 | 66.26 | PATRIC |
| Plasmids | 0 | 0 | PATRIC |
| Contig L50 | 9 | 8 | PATRIC |
| Genome Length | 5,445,390 bp | 6,509,961 bp | PATRIC |
| Contig N50 | 267,833 | 273,159 | PATRIC |
| Chromosomes | 0 | 0 | PATRIC |
| CDS | 4,951 | 6,239 | PATRIC |
| tRNA | 72 | 60 | PATRIC |
| Repeat Regions | 0 | 60 | PATRIC |
| rRNA | 3 | 7 | PATRIC |
| Hypothetical proteins | 2,087 | 1,307 | PATRIC |
| Proteins with functional assignments | 2,864 | 4,932 | PATRIC |
| Proteins with EC number assignments | 933 | 1,287 | PATRIC |
| Proteins with GO assignments | 806 | 1,091 | PATRIC |
| Proteins with Pathway assignments | 693 | 970 | PATRIC |
| Antibiotic Resistance | 0 | 51 | CARD |
| Antibiotic Resistance | 0 | 5 | NDARO |
| Antibiotic Resistance | 28 | 100 | PATRIC |
| Drug Target | 0 | 67 | DrugBank |
| Drug Target | 0 | 10 | TTD |
| Transporter | 2 | 186 | TCDB |
| Virulence Factor | 0 | 1 | PATRIC VF |
| Virulence Factor | 0 | 233 | VFDB |
| Virulence Factor | 0 | 86 | Victors |

The distribution of identified genes to cell subsystems is shown in Figure 1. *Sphingobacterium* S2 has more genes linked to protein processing, energy and membrane transport whereas *P. aeruginosa* S3 has more genes linked to metabolism, stress response/defense/virulence and cell envelop.

**Figure 1.**
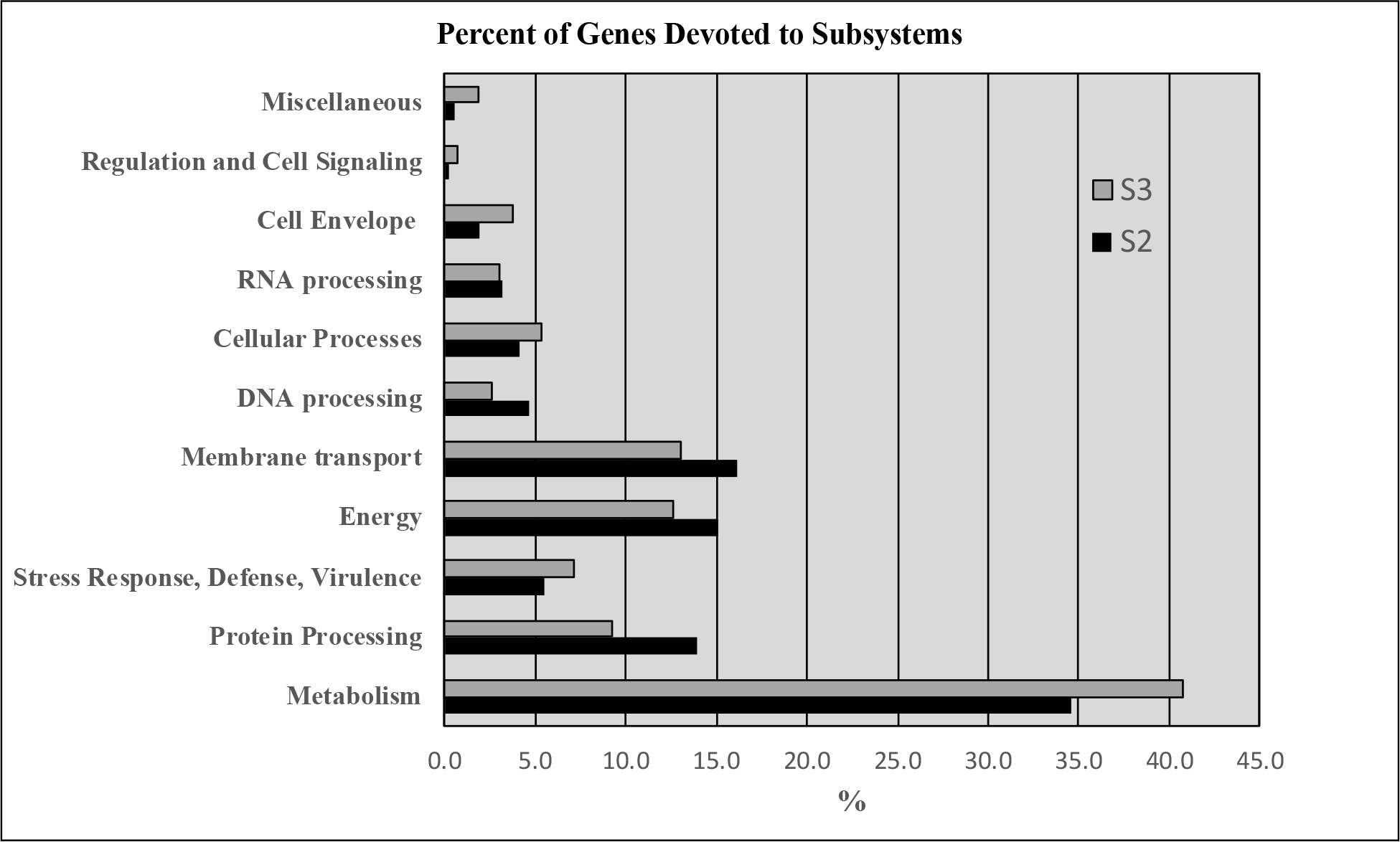
Distribution of identified genes to subsystems.

### 3.2 Genetic relatedness based on ANI

The average nucleotide identity (ANI) value describes the similarity between the sequences of the conserved regions of two genomes and measures the genetic relatedness between them [35]. ANI measurements are considered more informative over 16S rRNA gene identity as they are based on a larger number of genes [36]. ANI comparisons were done to explore the interspecies genetic relatedness between *Sphingobacterium sp*. strain S2 and *P. aeruginosa* strain S3 and other sequenced species from these genera. In the case of *P. aeruginosa* strain S3, Fig 1 shows a cluster was formed in the upper-right corner containing all the closely related strains with more than 98-99% gene identity based on 16S rRNA sequences and 93-97%. ANI. *P. aeruginosa* PSE305 showed highest ANI value and 16S rRNA gene identity of 97.69 % and 99 % respectively. Two or more strains with ANI values from 95% and above should be considered as the same species [37]. Accordingly, *P. aeruginosa* strain S3 and other strains included in the analysis belong to the same species. The results are consistent with by the phylogenetic results based on 16S rRNA gene identity as previously reported [12] but the ANI analysis indicated that our strain was closest to *P. aeruginosa* PSE305 rather than *P. aeruginosa* BUP2. Also, based on 16S rRNA comparative sequence analysis *P. aeruginosa* sv. O12 PA7 showed 99 % sequence similarity to *P. aeruginosa* strain S3 but had < 95 %. ANI value. ANI analysis for *Sphingobacterium sp*. strain S2 showed a different clustering of sequenced strains, some of which were not the same species. The data point in the upper-right corner represents *Sphingbacterium thalpophilum* DSM11723 which showed more than 98% 16S rRNA gene similarity and 98% ANI with Strain S2 and was the closest match to our strain. Two data points in the lower-right corner represented 97 and 98 % 16S rRNA gene identity but had ANI values of 79 and 80 % others having less than 95 % 16S rRNA gene identity had even lower ANI values. According to 16S rRNA gene identity and phylogenetic affiliations *Sphingobacterium sp*. strain S2 showed close resemblance to *Sphingobacterium thalpophilum* Y19 but ANI value showed *Sphingbacterium thalpophilum* DSM11723 to be the closest relative to our strain. Both of our *Sphingobacterium sp*. strain S2 and *P. aeruginosa* strain S3 were isolated from the compost samples, while the most closely related strains, *Sphingbacterium thalpophilum* DSM11723 and *P. aeruginosa* PSE305, were isolated from human clinical samples.

**Figure 2.**
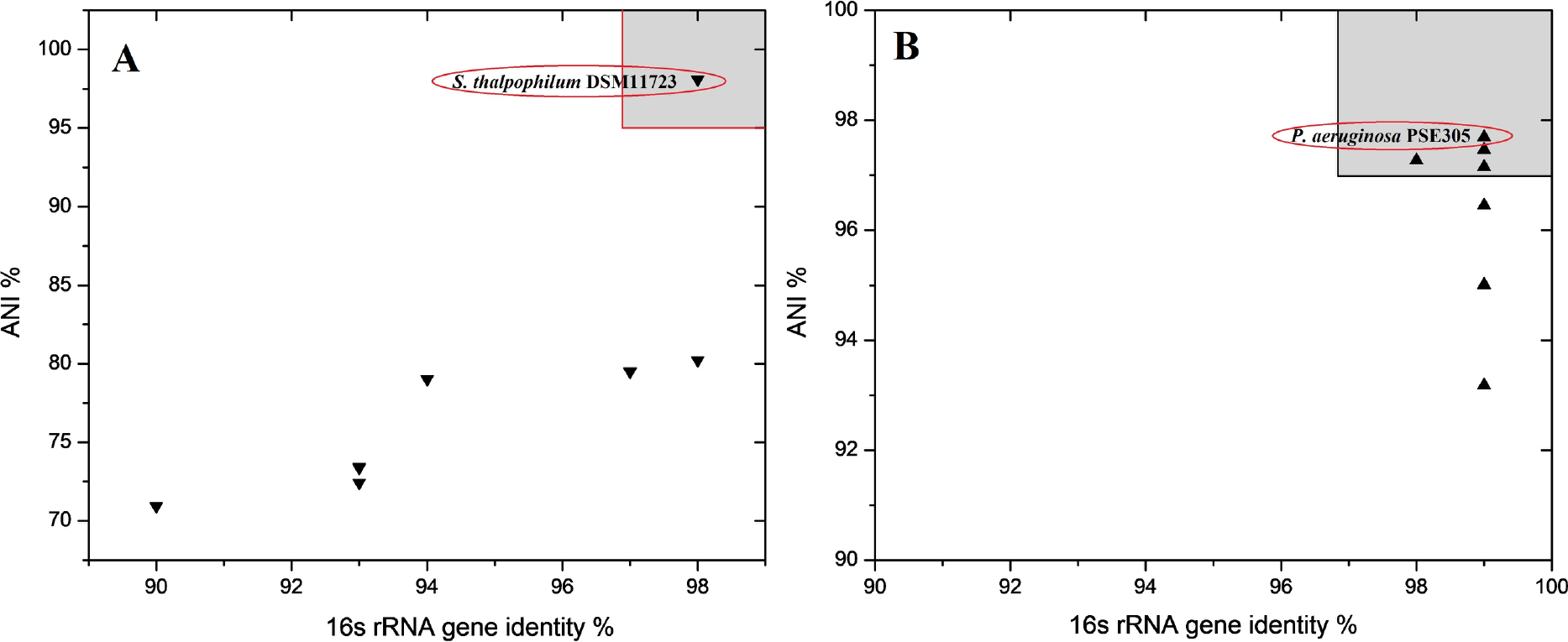
Correlation between 16S rRNA gene identity and ANI for pairs of genomes. (A) *Sphingobacterium sp* S2 and its closely related species, (B) *P. aeruginosa* S3 and its closely related species. Each triangle shows a relationship between the sequenced strains and one of its closely related species from the same genus. Reference in (A) *Sphingobacterium sp*. S2 (B) *P. aeruginosa* S3. Strains found within the grey area represents closely related species by both ANI and 16s rRNA.

### 3.3 MAUVE and MEDUSA alignments

MAUVE and MEDUSA were used to find sequence homologies of the two isolates with the strains identified as closest in the database by ANI calculations. The alignments shown in Figure 3 were derived by first aligning the SPADE-assembled contigs against reference strains, (*Sphingbacterium thalpophilum* DSM11723 or *P. aeruginosa* PSE305) with MeDuSa (REF), followed by alignment in MAUVE using the MeDuSa ordered scaffold of isolates S2 & S3. While the closest *Sphingobacterium* genome (DSM11723) in the current IMG database was close, based on a 98.1% ANI, this was a draft sequence with 32 contigs (Fig. 3, Panel A). As a result, the MAUVE alignment shows large regions of homology but spotty consensus in terms of synteny. In contrast, our *Pseudomonas* strain was identified as *P. aeruginosa* with many finished genomes of this species in the IMG database. ANI analysis (above) indicated that *P.aeruginosa* strain PSE305 was the closest. This strain has a completed genome and as a result, serves as an excellent reference strain for our isolate. Panel B of Figure 3 shows the putative alignment of *P.aeruginosa* S3 with *P. aeriginosa* PSE305. The 63 contigs of the SPADE assembled strain S3 was reduced to 9 contigs after alignment against PSE305 with MeDuSa.

**Figure 3.**
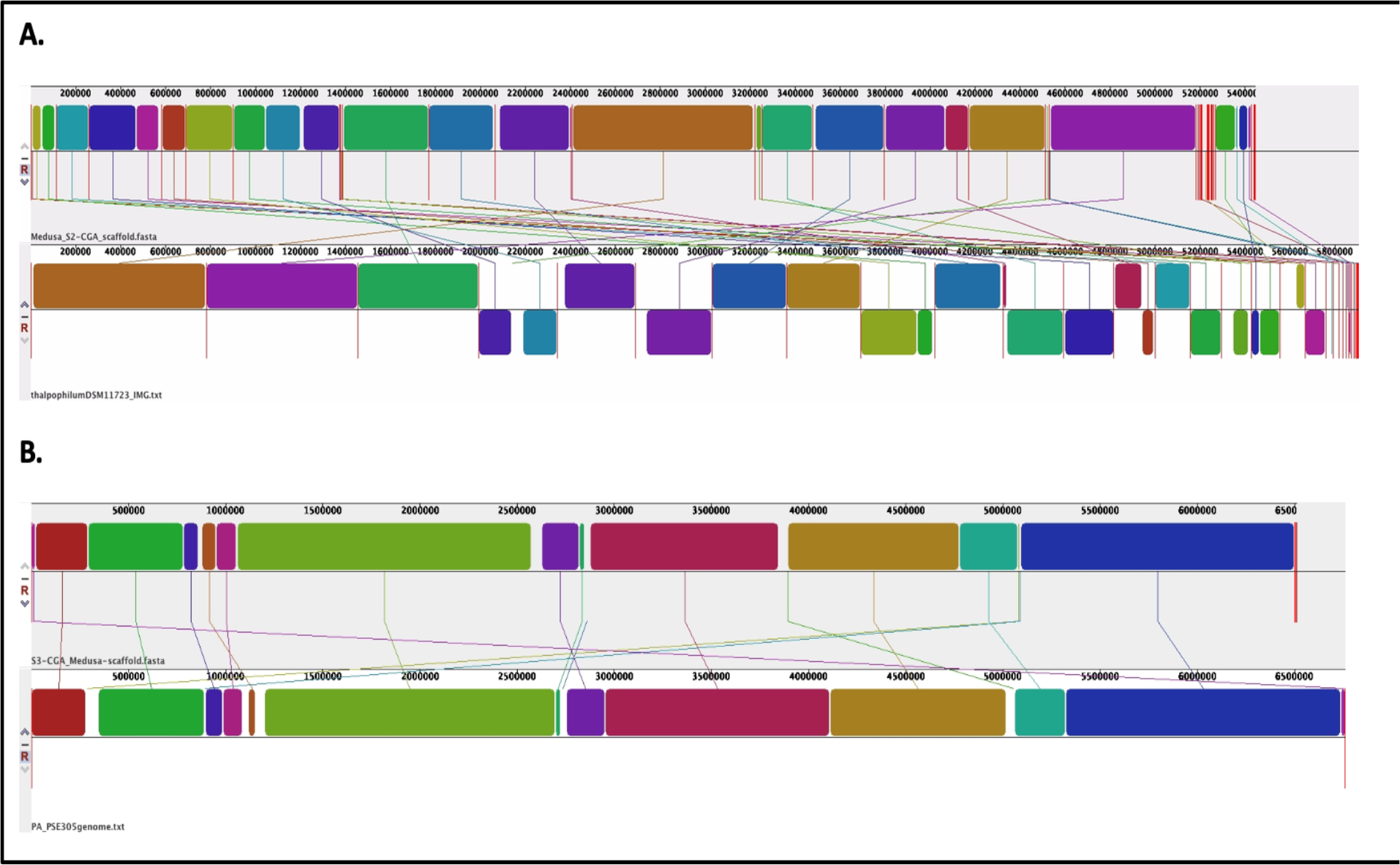
Genomic alignments generated in MeDuSa and MAUVE of *Sphingobacterium* sp. S2 with *Sphingbacterium thalpophilum DSM11723* (Panel A) and *P. aeruginosa* S3 (below) with *P. aeruginosa* PSE305 (Panel B).

### 3.4 Metabolism

*P. aeruginosa* is a Gram-negative bacterium able to grow aerobically and anaerobically, when nitrate is available as terminal electron acceptor. It is capable of thriving in highly diverse and unusual ecological niches with low availability of nutrients. Its metabolic versatility allows it to use a variety of diverse carbon sources including certain disinfectants [38]. Moreover, it can synthesize a number of antimicrobial compounds [24, 25]. Using genome annotation through PATRIC, 1,084 ORFs in *P.aeruginosa* were assigned to metabolic pathways. Among the major pathways, 361 genes were assigned to amino acid metabolism had 361 genes assigned to it and 192, 286, 225 and 168 genes were assigned for energy metabolism, carbohydrate metabolism, metabolism of cofactors and vitamins and lipid metabolism, respectively. Metabolism of xenobiotics was allocated to 165 genes while 129 and 118 genes were nucleotide metabolism and biosynthesis of secondary metabolites had 129 and 118 genes assigned to their functions.

*Sphingobacterium spp*. are gram-negative rods, aerobic, exhibiting sliding motility and form yellow-pigmented colonies. *Sphingobacteria* have been isolated from diverse environments like soil, water, compost, deserts, blood and urine samples from human patients. A distinctive feature of *Sphingobacterum* is the presence of sphingolipids in their cell wall in high concentrations [14, 39]. In *Sphingobacteria S2*, 820 ORFs were assigned metabolic functions. 212 genes were identified as participating in amino acid metabolism while 262 genes were dedicated to carbohydrate metabolism. Energy metabolism and lipid metabolism had 148 and 132 genes identified, respectively, while 67 genes were assigned to xenobiotic biodegradation and metabolism. *The disparity in assigned genes between Sphingobacterium and P. aeruginosa, particularly with respect to carbohydrate metabolism and amino acid synthesis* is presented in Figure 4.

**Figure 4.**
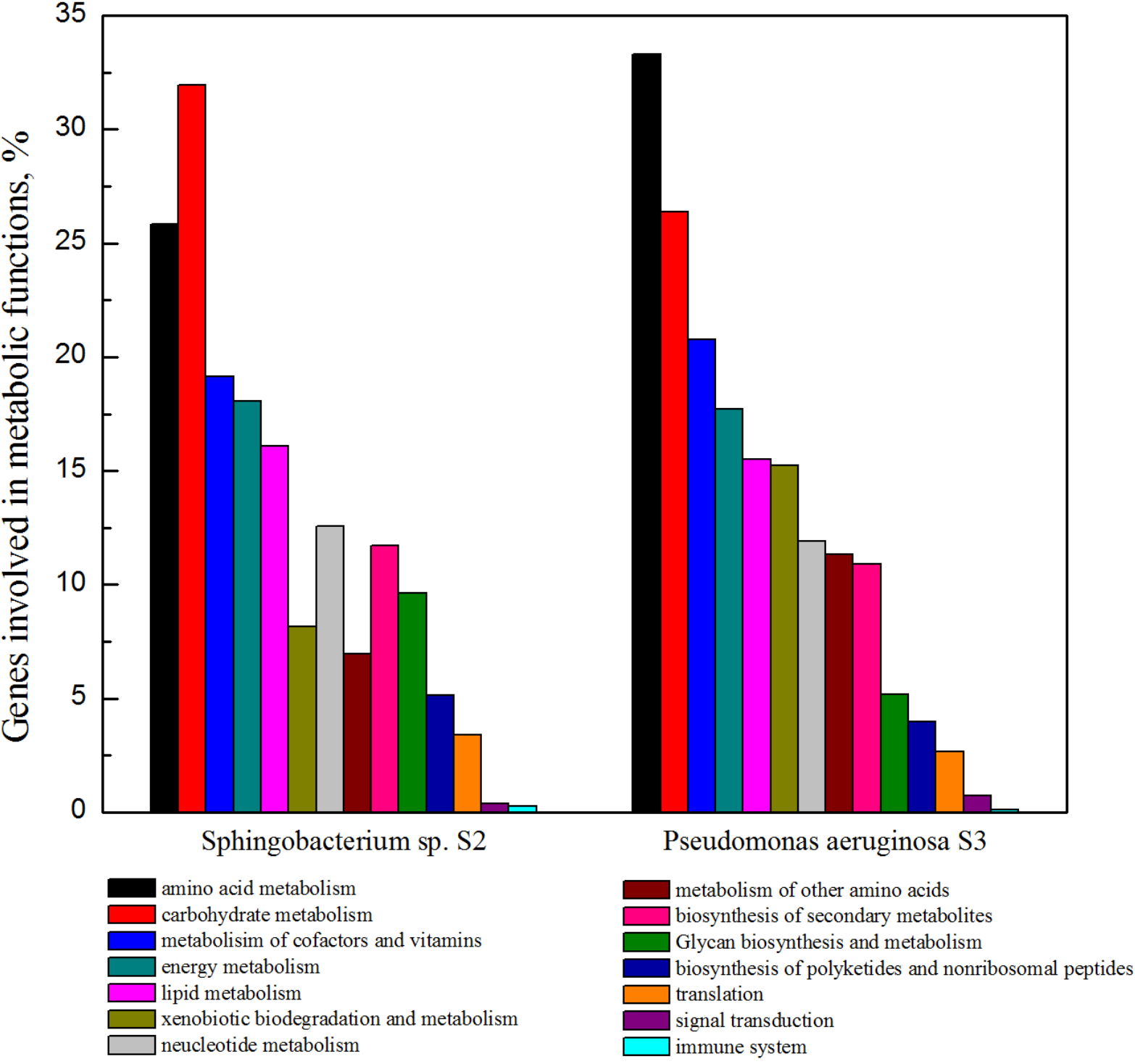
Metabolic features of *P. aeruginosa* S3 and *Sphingobacterium sp*. S2

### 3.3. Xenobiotics biodegradation metabolism

In previous reports the role of *P. aeruginosa* in degradation of different polymers including PAHs, biodegradation of xenobiotic compounds, degradation of oil, dyes and plastics as well is well documented [26–30], *Sphingobacterium* had also been reported to have their potential role in biodegradation of different pollutants including mixed plastic waste, PAHs, biodegradation of oil and pesticides [19–21]. Table 2 describes the major pathways and the number of genes related to biodegradation of different xenobiotic compounds in both strains S2 and S3. According to sequence analysis, several pathways for the biodegradation of xenobiotic compounds were found in both strains with twice as many detected in *P. aeruginosa* as compared to *Sphingobacterium sp. This may be due to the fact that P. aeruginosa* is more comprehensively studied compared to *Sphingobacterium sp*. Among all the degradation pathways found in both strains, the degradation of benzoate in *P. aeruginosa had the greatest number of annotated genes* Benzoate, an aromatic compound, has been widely used as a model for the study of the bacterial catabolism of aromatic compounds. In *Sphingobacterium sp*. genes associated with 1,4-Dichlorobenzene degradation were detected and this compound is a well-studied halogenated aromatic hydrocarbon [40]. Many genes related to the degradation pathway of one of the most important class of pollutants, polycyclic aromatic hydrocarbons (PAHs) like naphthalene, anthracene, 1- and 2-methylnaphthalene, was found in both strains. Among the halogenated organic compounds, 29 genes were dedicated to tetrachloroethene degradation in *P. aeruginosa S3*, while 6 genes were found in *Sphingobacterium sp. S2*. For aromatic compounds and chlorinated aromatic compounds, pathways for the biodegradation of toluene, trinitrotoluene, xylene degradation, 1,4-Dichlorobenzene degradation and 2,4-Dichlorobenzoate were also found. Genes for the biodegradation of Bisphenol A, one of the most abundantly produced chemicals released into the environment and is a serious environmental pollutant, was also found in both of our strains. *P. aeruginosa* S3 was found to have 13 genes for bisphenol A degradation pathway[41]. Pathways for pesticides degradation like 1,1,1-Trichloro-2,2-bis(4-chlorophenyl) ethane (DDT) degradation and atrazine biodegradation were also found in both the isolates.

**Table 2.** Pathways and number of genes involved in aromatic compound metabolism in draft genome sequence of *P. aeruginosa S3* and *Sphingobacterium sp. S2*. Unique EC count refers to the number of unique enzymes identified.

|  | *Pseudomonas aeruginosa* (S3) |  | *Sphingobacterium* sp. (S2) |  |
| --- | --- | --- | --- | --- |
| Pathway Name | Unique Gene Count | Unique EC Count | Unique Gene Count | Unique EC Count |
| 1- and 2-Methylnaphthalene degradation | 20 | 8 | 8 | 6 |
| 1,4-Dichlorobenzene degradation | 43 | 13 | 18 | 6 |
| 2,4-Dichlorobenzoate degradation | 17 | 9 | 7 | 3 |
| Atrazine degradation | 10 | 2 | 7 | 2 |
| Benzoate degradation via hydroxylation | 60 | 25 | 16 | 7 |
| Biphenyl degradation | 3 | 2 | 3 | 2 |
| Bisphenol A degradation | 13 | 6 | 4 | 3 |
| Caprolactam degradation | 21 | 6 | 9 | 4 |
| Drug metabolism - cytochrome P450 | 19 | 3 | 3 | 2 |
| Drug metabolism - other enzymes | 9 | 9 | 8 | 8 |
| Ethylbenzene degradation | 10 | 3 | 6 | 3 |
| Fluorobenzoate degradation | 9 | 7 | 1 | 1 |
| γ-Hexachlorocyclohexane degradation | 9 | 6 | 2 | 2 |
| Geraniol degradation | 30 | 9 | 9 | 4 |
| Naphthalene and anthracene degradation | 16 | 7 | 6 | 3 |
| Styrene degradation | 14 | 7 | 2 | 2 |
| Tetrachloroethene degradation | 29 | 8 | 6 | 4 |
| Toluene and xylene degradation | 14 | 6 |  | 2 |
| Trinitrotoluene degradation | 12 | 4 | 10 | 3 |
| DDT degradation | 5 | 4 | 2 | 2 |
| Metabolism of xenobiotics by cytochrome P450 | 19 | 3 | 5 | 3 |

### 3.4. Lactate metabolism

Lactate utilization as sole carbon source is a property of many bacteria, where a key step of the process is oxidation of lactate [42–46]. Lactate dehydrogenases found in microbes are of two types, NAD-dependent lactate dehydrogenases (nLDHs) and NAD-independent lactate dehydrogenases (iLDHs), also called respiratory lactate. The latter is usually considered to be the enzyme mainly responsible for metabolism of lactate as a carbon source [45]. The lactate utilization system is comprised of three main membrane bound proteins: NAD-independent L-lactate dehydrogenase (L-iLDH), NAD-independent D-lactate dehydrogenase (D-iLDH), and a lactate permease (LldP). Lactate permease, LldP is responsible to take up lactate into the cells and lactate dehydrogenases carry out the oxidation of either form of lactate to pyruvate [47, 48]. In pathogenesis of some microbes role of lactate utilization has been observed [46]. Lactate utilization in pathogens not only has stimulatory effect on their growth, it also enhances synthesis of pathogenic determinants and imparts them resistance against various bactericidal mechanisms [46]. Utilization of lactate by different *Pseudomonas* strains is very well documented [44, 49–51]. In sequence analysis of our *P. aeruginosa* strain S3, a complete cascade of genes was found, encoding the machinery for lactate utilization that included a L-lactate permease, both L-lactate dehydrogenase and D-lactate dehydrogenase and a Lactate-responsive regulator LldR Table 3. This strain was isolated and characterized for its potential to degrade Poly(lactic acid), one of the more promising of the bio-based and biodegradable polymers currently in the market, and its potential to utilize lactate, one of the final products of PLA degradation, as a sole carbon source was already established in our previous study [12]. Presence of the lactate utilization machinery found through genome sequencing is the confirmation of our previous findings regarding lactate utilization by *P. aeruginosa*, strain S3. It was previously described in the literature that both L-iLDH and D-iLDH are present in the single operon and are induced coordinately in all the reported *Pseudomonas* strains. Expression of both enzymes is controlled by the presence of enantiomer of lactate [48]. In previous studies, it was reported that in *P. aeruginosa* strain XMG lactate utilization operon lldPDE consists of genes for lactate permease LldP, the L-lactate dehydrogenase LldD and the D-lactate dehydrogenase LldE and a nearby located lldR gene, which codes for a regulator LldR [48, 52]. In the genome analysis of our isolate *Sphingobacterium* strain S2, we found an incomplete set of genes, both L-lactate dehydrogenase and D-lactate dehydrogenase were present but no lactate permease was detected Table 3. Absence of lactate permease suggested that the strain is incapable of utilizing lactic acid as carbon source. This again confirmed our previous findings that showed that *Sphingobacterium* strain S2 did not utilize lactic acid as sole source of carbon. *Sphingobacterium* strain S2 was isolated and characterized based on its ability to degrade PLA suggesting that another degradation product of PLA was utilized for growth. Inability to grow on lactic acid had been previously reported in literature for different strains of *Sphingobacterium* [18, 53].

**Table 3.** Genes for lactate metabolism in *P. aeruginosa* S3 and *Sphingobacterium sp*. S2

| **Proteins for lactate utilization in *Pseudomonas aeruginosa* S3** |  |  |
| --- | --- | --- |
| **Description** | **AA length** | **Proteins** |
| L-lactate permease | 560 | 1 |
| L-lactate dehydrogenase | 383 | 2 |
| D-lactate dehydratase (EC 4.2.1.130) | 291 | 1 |
| D-lactate dehydrogenase (EC 1.1.1.28) | 329 | 1 |
| Acetolactate synthase large subunit (EC 2.2.1.6) | 574 | 1 |
| Acetolactate synthase small subunit (EC 2.2.1.6) | 163 | 2 |
| Lactate-responsive regulator LldR, GntR family | 257 | 1 |
| Predicted D-lactate dehydrog., Fe-S protein, FAD/FMN-containing | 938 | 1 |
| **Proteins for lactate utilization in *Sphingobacterium* sp. S2** |  |  |
| **Description** | **AA length** | **Proteins** |
| L-lactate dehydrogenase | 389 | 1 |
| D-lactate dehydratase (EC 4.2.1.130) | 145 | 2 |
| D-lactate dehydrogenase (EC 1.1.1.28) | 330 | 1 |
| Acetolactate synthase large subunit (EC 2.2.1.6) | 606 | 1 |
| Acetolactate synthase small subunit (EC 2.2.1.6) | 196 | 1 |
| Fe-S protein, homolog of lactate dehydrogenase SO1521 | 974 | 1 |
| Predicted L-lactate dehydrogenase, Fe-S oxidoreductase subunit YkgE | 242 | 1 |
| Predicted L-lactate dehydrogenase, hypothetical protein subunit YkgG | 215 | 1 |
| Predicted L-lactate dehydrogenase, Iron-sulfur cluster-binding subunit YkgF | 462 | 1 |

### 3.5. Genetic determinants for biofilm formation and regulation

In contrast to the planktonic life style, cells within a biofilm matrix are in close proximity where secreted enzymes provide optimal returns for the population [54]. The phenomenon of microbial biofilm formation is also related to other survival strategies like metal and antimicrobial resistance, tolerance and bioremediation [55, 56]. Application of biofilm mediated bioremediation has been found superior to other bioremediation strategies and is being applied in bioremediation of different environmental pollutants [57–61]. Microorganisms that develop a biofilm and have the ability to secrete polymers establishing a protective extracellular matrix, are physiologically more resilient to environmental changes, making them a logical choice for the treatment of different pollutants. These microbes use different strategies like biosorption, bioaccumulation and biomineralization to slowly degrade compounds [62].

Biofilm formation by Pseudomonas species had been well documented in literature[63]. *P. aeruginosa* is a remarkably adept opportunist with striking ability to develop biofilm [64]. In our previous study, we also observed biofilm formation by our isolate *P. aeruginosa* strain S3 on the surface of PLA during the process of biodegradation [12]. Phenomenon of biofilm formation on the surface of PLA had previously been reported by other authors as well [65–67].

In the genetic analysis of our isolate we found factors involved in the development of matrix of *P. aeruginosa* biofilm and its regulation (Table 4). The genes for three types of exopolysaccharides (EPS), previously reported as involved in construction of biofilm matrix of *P. aeruginosa*, (Pel, Psl and alginate [64, 68]) were found in our isolate. These EPS molecules form the protective matrix [69]. Psl is the primary factor in charge of the initiation and maintenance of biofilm structure by providing cell to cell and cell to surface interactions [70–73]. It also works as a signaling molecule to the successive events involved in the formation of biofilms and also acts as a defensive layer for different immune and antibiotic attacks [74]. Pel polysaccharide is a glucose-rich extracellular matrix and is involved in the formation of biofilms that are attached to the solid surfaces. It is considered to be less important compared to Psl [64, 73, 75, 76]. In *P. aeruginosa* from clinical isolates of CF patients Alginate is produced [23]. Besides its role in maintenance and protection of biofilm structure, it is essential for water and nutrient preservation [77].

**Table 4.** Genetic elements detected in *P. aeruginosa* S3 and previously identified as involved in biofilm formation in other strains of *P. aeruginosa*

| **Factors involved in Biofilm formation and regulation detected in *P. aeruginosa* S3** |  |
| --- | --- |
| **Psl polysaccharide** | **Pseudomonas quinolone signal (PQS)** |
| Extracellular Matrix protein PslA | PQS biosynthesis protein PqsH, similar to FAD-dependent monooxygenases |
| Extracellular Matrix protein PslC | PQS biosynthesis protein PqsA, anthranilate-CoA ligase (EC 6.2.1.32) |
| Extracellular Matrix protein PslD | PQS biosynthesis protein PqsB, similar to 3-oxoacyl-[acyl-carrier-protein] synthase III |
| Extracellular Matrix protein PslE | PQS biosynthesis protein PqsC, similar to 3-oxoacyl-[acyl-carrier-protein] synthase III |
| Extracellular Matrix protein PslF | PQS biosynthesis protein PqsD, similar to 3-oxoacyl-[acyl-carrier-protein] synthase III |
| Extracellular Matrix protein PslG | PqsE, quinolone signal response protein |
| Extracellular Matrix protein PslL | Anthranilate synthase, aminase component (EC 4.1.3.27) |
| Extracellular Matrix protein PslJ | Anthranilate synthase, amidotransferase component (EC 4.1.3.27) |
| Extracellular Matrix protein PslK | Multiple virulence factor regulator MvfR/PqsR |
|  | Putative transcriptional regulator near PqsH |
| **Pel Polysaccharide** | **c-di-GMP** |
| Extracellular Matrix protein PelG | 3',5'-cyclic-nucleotide phosphodiesterase (EC 3.1.4.17) |
| Extracellular matrix protein PelF, glycosyltransferase, group 1 | 5'-nucleotidase/2',3'-cyclic phosphodiesterase and related esterases |
| Extracellular Matrix protein PelE | Acyl carrier protein phosphodiesterase (EC 3.1.4.14) |
| Extracellular Matrix protein PelD | diguanylate cyclase/phosphodiesterase (GGDEF & EAL domains) with PAS/PAC sensor(s) |
| Extracellular Matrix protein PelC | Glycerophosphoryl diester phosphodiesterase (EC 3.1.4.46) |
| Extracellular Matrix protein PelB | Phosphodiesterase/alkaline phosphatase D |
| Extracellular Matrix protein PelA | Membrane bound c-di-GMP receptor LapD |
| **Alginate** | **Quorum sensing systems** |
| Alginate regulatory protein AlgQ | N-3-oxododecanoyl-L-homoserine lactone quorum-sensing transcriptional activator @ Transcriptional |
| Alginate regulatory protein AlgP | regulator LasR |
| Alginate biosynthesis protein AlgZ/FimS | N-acyl-L-homoserine lactone synthetase LasI |
| Alginate biosynthesis transcriptional regulatory protein algB | N-butyryl-L-homoserine lactone quorum-sensing transcriptional activator @ Transcriptional |
| Alginate biosynthesis protein Alg8 | regulator RhIR |
| Alginate biosynthesis protein Alg44 |  |
| Alginate biosynthesis protein AlgK precursor |  |
| Outer membrane protein AlgE |  |
| Alginate biosynthesis protein AlgX |  |
| Alginate lyase precursor (EC 4.2.2.3) |  |
| Alginate biosynthesis protein AlgJ |  |
| Alginate o-acetyltransferase AlgF |  |
| Alginate biosynthesis transcriptional activator |  |

Biofilm formation is a multicellular process stimulated by both environmental signals controlled by regulatory networks. During the biofilm formation, cells undergo many phenotypic shifts that are regulated by a large array of genes [78]. In the genome of *P. aeruginosa* strain S3, several regulatory factors were identified. One of these regulatory factors was the signaling molecule bis-(3′-5′)-cyclic dimeric guanosine monophosphate (c-di-GMP) which is considered as one of the most significant molecular determinants in biofilm regulation [79]. A c-di-GMP molecule controls the interchange between the planktonic and biofilm-associated lifestyle of bacteria by stimulating the adhesins biosynthesis and exopolysaccharides during formation of biofilm [80]. The bacterial cell to cell communication system known as quorum sensing (QS) is involved in the maintenance of many biological processes like biofilm formation, bioluminescence, antibiotics production, virulence factor expression, competence for DNA uptake, and sporulation [81, 82].

LasR/LasI, RhlR/RhlI and PQS are the three quorum sensing signaling systems employed by *P. aeruginosa* to control biofilm formation [83, 84]. These three QS signaling system were found to be the part of the genome for our isolate.

There is little information in the literature regarding the genetic elements involved in biofilm formation in *Sphingobacterium*, Genome analysis of our isolate *Sphingobacterium sp*., strain S2 showed presence of genes for Stage 0 sporulation protein YaaT. This protein is reported to be involved in the sporulation process and biofilm development [85, 86].

### 3.6. Enzymes

Biodegradation of polymers is carried out by two types of enzymes, extracellular enzymes that degrade long chain polymers into short oligomers or subunits that are subsequently carried inside the cell, and intracellular enzymes that further degrade the small transported units [87, 88]. Degradation of synthetic polymers in the environment can be a slow process [89, 90]. PLA is a synthetic linear aliphatic polyester of lactic acid monomers joined together by ester linkages [3]. The presence of ester bonds in its backbone make the polymer sensitive to hydrolysis, both chemically as well as enzymatically [88]. Biodegradation of polyesters is mostly carried out by esterolytic enzymes such as esterases, lipases, or proteases. Microbial carboxyl esterases: classification, properties and application in biocatalysis). In the literature microbial degradation of PLA is mainly reported by proteases, lipses, esterses and a few cutinases as well [4]. Both *Sphingobacterium sp*. and *P. aeruginosa* had been documented before to have a role in the degradation of different environmental pollutants such as mixed plastic waste, PAHs, oil, and dyes and pesticides, *P. aeruginosa* has also been reported to have potential to degrade PLA nano-composites [19, 26, 28, 91, 92]. In our previous study we characterized the degradation of PLA by our isolates [12]. Genome annotation of of both strains report presence of hydrolytic enzymes in their genomes, putatively related to their degradation of PLA (Table 5). In the genome of *P. aeruginosa* S3, 75 different types of proteases, 50 esterases and 25 different types of lipases were detected.

**Table 5.** Different type of hydrolytic enzymes found in the draft genome of *P. aeruginosa* S3 and *Sphingobacterium sp*. S2

| Type of enzymes | *Pseudomonas aeruginosa* S3 |  | *Sphingobacterium sp.* S2 |  |
| --- | --- | --- | --- | --- |
|  | Common to reference genomes | Unique to Strain S3 | Common to reference genomes | Unique to Strain S2 |
| **Hydrolase** | **123** | **51** | **84** | **8** |
| **Lipase** | **20** | **5** | **10** | **8** |
| **Protease** | **57** | **18** | **22** | **14** |
| **Esterase** | **33** | **17** | **25** | **5** |
| **Phosphodiesterase** | **8** | **5** | **5** | **4** |
| **Oxygenase** | **63** | **19** | **7** | **3** |
| **Catalase** | **6** | **6** | **2** | **1** |
| **Phosphatase** | **66** | **17** | **25** | **13** |

Similarly, in the genome of *Sphingobacterium sp*. S2 36 proteases, 30 esterases and 18 lipases were identified. Apart from these various phosphodiesterases, oxygenases, catalases and phosphatases have also been found in the genome of both isolates. Already established potential of these strains to degrade various environmental pollutants is reaffirmed by the presence of these diverse enzymes.

## 4. Conclusion

The present study reports the whole genome sequence analysis of two bacterial strains *P. aeruginosa* S3 and *sphinogobacterium sp*. S2, isolated from the compost and having the potential to degrade poly(lactic acid), PLA, at mesophillic temperatures (~30°C). Draft genomes of both the strains were studied to gain an insight into the genetic elements that are involved in conferring different properties to the strains helping in the degradation of PLA. The catabolic genes responsible for biodegradation of different xenobiotic compounds, genes responsible for formation and regulation of biofilm, genes for transport and utilization of lactate and several enzymes predicted to be involved in the degradation of many organic pollutants were identified on the graft genomes. All these characteristics demonstrate the degradation potential of the strains against PLA observed in the previous studies by our group; importantly it gives insight into the possible enzymes involved in the degradation of the polymer.

## 5. Acknowledgements

The authors show our gratitude to the Higher Education Commission of Pakistan (HEC) and the Government of Pakistan for providing a fellowship for S.S. through the international research support initiative program (IRSIP). We are also thankful to The Genomics Facility (MSU-RTSF) at Michigan State University (East Lansing, MI, USA) for performing whole genome sequencing for our strains.

